# Thiamine availability and acquisition differ between natural and controlled environments

**DOI:** 10.64898/2026.06.04.730186

**Authors:** Matthew H. Futia, Colin Clark, Christopher P. Suffridge, Gillian St. John, J. Ellen Marsden, Jacques Rinchard

## Abstract

Thiamine Deficiency Complex (TDC) is a reproductive disorder that affects recruitment of diverse salmonine populations globally. Typical symptoms include behavioral and neurological abnormalities and high offspring mortality. TDC is common in hatcheries that rear salmonines obtained from wild populations, and symptoms are mitigated by thiamine treatment. However, no studies have quantified thiamine concentrations in wild embryos. Here, we evaluated whether fertilized eggs and/or embryos may acquire thiamine from natural sources (e.g., biotic breakdown products and diet) during development. Lake trout (*Salvelinus namaycush*) gametes were obtained from feral adults in Lake Champlain and fertilized eggs were grouped by family with paired rearing under natural (Lake Champlain) and artificial (controlled laboratory) conditions. Average thiamine concentrations were similar between lake-reared and laboratory-reared fish prior to hatch; however, lake-reared fish experienced significant increases in thiamine concentrations at and after hatching compared to previous stages and compared to laboratory-reared fish; laboratory-reared fish experienced no increases in thiamine concentrations. Water samples revealed an abundance of thiamine precursors and byproducts in the natural environment, which may serve as sources of thiamine for developing embryos. These results demonstrate that salmonine embryos can acquire thiamine from natural sources during development, which may mitigate effects of TDC.

## Introduction

Thiamine Deficiency Complex (TDC) is a natural phenomenon associated with a vitamin B_1_ (thiamin(e)) deficiency that affects salmonine populations globally^[1]^. The deficiency is often caused by disruptions in the food web and results in adult females allocating reduced concentrations of thiamine to their oocytes and producing eggs with low thiamine concentrations^[2]^. When egg thiamine concentrations are low, post-hatch free embryos can experience behavioral and physiological issues (e.g., abnormal swimming, lethargy, reduced immune response) that cause direct mortality in severe cases^[3,4]^. In addition, free- and post-embryo behaviors including foraging efficiency and predator avoidance can be impaired by thiamine deficiency, resulting in additional offspring mortality^[5,6]^. Due to the elevated offspring mortality associated with TDC, this syndrome has been identified as a major threat to salmonine conservation and restoration efforts^[1,7]^.

Lake trout (*Salvelinus namaycush*) recruitment has been low in multiple lakes throughout the Laurentian Great Lakes (hereafter referred to as Great Lakes) region for many decades, and successful restoration of their wild populations has been limited to a few locations^[8–10]^. Multiple potential impediments have been identified (e.g., sea lamprey predation, overfishing, habitat alterations), including TDC^[11]^. Symptoms of TDC in the Great Lakes region were first documented in 1968 and included elevated offspring mortality in hatcheries ^[12]^, although TDC may have occurred much earlier in some parts of the basin^[13]^. Nearly 20 years later, thiamine deficiencies were shown to cause the elevated offspring morality^[4,14]^ and since then the deficiency has been linked to consumption of invasive alewife (*Alosa pseudoharengus*)^[15]^. For example, while alewife were the primary prey for lake trout in Lake Huron, lake trout recruitment was minimal; following the collapse of the alewife population in the 2000s, lake trout recruitment increased substantially presumably due to relief from TDC^[16,17]^. However, lake trout recruitment has been observed in lakes where alewife are consumed by lake trout. In Keuka Lake (New York, USA), alewife and lake trout coexisted for many years without limiting wild lake trout recruitment^[18]^. In Lake Champlain (Vermont and New York, USA), lake trout recruitment first appeared after alewife became established in the lake^[10]^, resulting in complete population recovery (i.e., termination of stocking programs) in 13 years^[19]^ (M. Murphy, VTFWD, pers. comm.) despite low thiamine levels in lake trout and Atlantic salmon (*Salmo salar*)^[20]^.

Although the effects of TDC have been studied for decades, elevated offspring mortality caused by TDC has only been studied in laboratory and hatchery settings^[12,14,21]^ and has yet to be evaluated in natural conditions. However, observing thiamine-deficient free and post embryos in the wild is challenging considering direct observation is impractical and capturing thiamine-deficient embryos in fry traps is unlikely due to their inability to swim into the traps. Therefore, the lack of documented TDC in wild embryos could be due to the inability to detect those individuals. An additional hypothesis for lack of TDC in wild embryos is that eggs and/or embryos may acquire thiamine from their surrounding environment, thereby reducing or even eliminating TDC under natural conditions. In hatchery settings, thiamine treatments have been successfully used at various life stages to mitigate the impacts of TDC^[22–24]^. Exposing eggs to thiamine-enriched water baths (≥ 1,000 mg/L) during water hardening resulted in significantly reduced TDC-induced offspring mortality^[22,23]^. Similarly, immersing free embryos in thiamine baths or at the swim-up stage lowered TDC-related mortality^[24]^. These studies demonstrate that both eggs and free embryos can acquire thiamine from the surrounding water when thiamine is present in high concentrations.

In natural environments (i.e., spawning reefs), it is likely that thiamine autotrophs and decomposing biota enrich the surrounding water with thiamine. If thiamine is present in the water, developing embryos may be able to uptake sufficient concentrations during the five months of incubation and potentially offset TDC naturally. Recently, analytical developments have enabled quantification of dissolved thiamine-related compounds (dTRC), likely microbially produced, in natural environments^[25]^. Therefore, it is now possible to test whether dTRCs in ambient water are sufficient to increase embryo thiamine concentrations overwinter. Lake trout free- and post-embryos may also be able to acquire ambient thiamine by ingestion after hatching. In the wild, free embryos begin to feed on zooplankton prior to yolk-sac depletion, within two weeks of hatching^[26]^; because zooplankton are rich in thiamine^[27]^, early feeding may be able to mitigate the impacts of TDC by providing thiamine supplementation. Monitoring dTRC concentrations in water collected from rearing environments throughout embryo development as well as concentrations in eggs and free embryos under natural and controlled laboratory conditions can be used to demonstrate whether acquisition of ambient thiamine occurs during development. Whether thiamine uptake during development alleviates TDC in the wild can then be evaluated by comparing free-embryo feeding success and survival to thiamine concentrations in eggs and embryos.

Here, we conducted the first study to evaluate thiamine acquisition by salmonine embryos reared in a natural environment. The study took place in Lake Champlain where TDC has been observed in native salmonine populations, including lake trout^[20]^. Eggs from individual wild-caught females were split into two groups and reared in either a natural setting in Lake Champlain (lake group) or in laboratory conditions (lab group). Eggs or embryos were sampled at four periods during development (unfertilized eggs, fertilized eggs, free embryos at hatch, and free embryos post hatch) to determine changes in tissue thiamine concentrations and relative abundance of thiamine forms (i.e., vitamers) that were dependent on rearing environment. Tissue thiamine concentrations were measured using dehydrated samples (hereafter referred to as dry thiamine), which accounted for changes in water content among sampling periods, in addition to wet samples (hereafter referred to as wet thiamine) that are typically analysed^[28]^. We also sampled for ambient sources of thiamine (i.e., dTRC and zooplankton in stomachs) available to embryos in their rearing environment. Water samples were collected at a second site in Lake Champlain to provide additional evaluation of dTRC availability in lake water. The purpose of this project was to establish whether embryos with low thiamine concentrations can acquire ambient thiamine during egg development or at the free embryo stage post hatch in natural settings. Such information may have critical management implications if embryos developing in natural environments can acquire enough thiamine to offset mortality associated with TDC.

## Results

### Sample acquisition and developmental stages

Sampling of free embryos at hatch for the lake-reared group occurred as soon as the field site was accessible after ice-melt on Lake Champlain. Four of the 20 families had live unhatched eggs remaining at this sampling period, but these were limited to one to three eggs per family. Only two egg bags had complete mortality prior to hatch (i.e., only dead eggs recovered) including one at hatch and one at the final sampling period approximately five weeks later; all other egg bags had at least one free embryo (dead or alive) present. Dead embryos were typically more common than live embryos, but many embryos appeared freshly dead (minimal discoloration). Multiple free embryos were also observed escaping from egg bags while removing the bags during both free-embryo sampling periods, limiting accurate counts of live embryos. Free embryos collected from the lake at the final sampling interval were found at a range of sizes, from 22 to 28 mm, and yolk sacs were nearly completely absorbed. Lab free embryos were sampled daily at hatch for 18 families; for the other two families, all embryos died prior to hatch or there were insufficient remaining embryos to measure thiamine. Final sampling of free embryos reared in the lab occurred for 19 families approximately three months after hatch.

### Thiamine acquisition

Total thiamine (hereafter referred to as thiamine) concentrations in unfertilized eggs ranged between 4.05 and 12.90 nmol/g among the 20 females in this study (Supplementary Table 1). Compared to those initial concentrations, wet thiamine concentrations increased significantly in lake-reared embryos while concentrations decreased over time in the lab group (Figure 1). In the lake, wet thiamine concentrations initially decreased significantly from unfertilized eggs to fertilized embryos (p = 0.001; estimated marginal means) by an average of 30%. Embryo thiamine concentrations then increased by 120% between fertilization and hatch and were significantly greater at hatch compared to unfertilized and fertilized samples (p < 0.001; estimated marginal means). Average thiamine concentrations in embryos from the lake group further increased by 27% between hatch and the final sampling of free embryos, but this change was not significant (p = 0.918; estimated marginal means). Eggs reared in the lab also experienced a significant decline in thiamine concentrations following fertilization compared to unfertilized eggs (p < 0.001; estimated marginal means) with an average decline of 48%, but concentrations remained low and consistent in the following sampling periods (p > 0.729; estimated marginal means; Figure 1). Pairwise comparisons between lake- and lab-reared embryos demonstrated significantly greater wet thiamine concentrations in lake-reared embryos following fertilization (average difference of 30%; p = 0.018; estimated marginal means), at hatch (average difference of 112%; p < 0.001; estimated marginal means), and at the final sampling period (average difference of 122%; p < 0.001; estimated marginal means; Figure 1).

**Figure 1.**
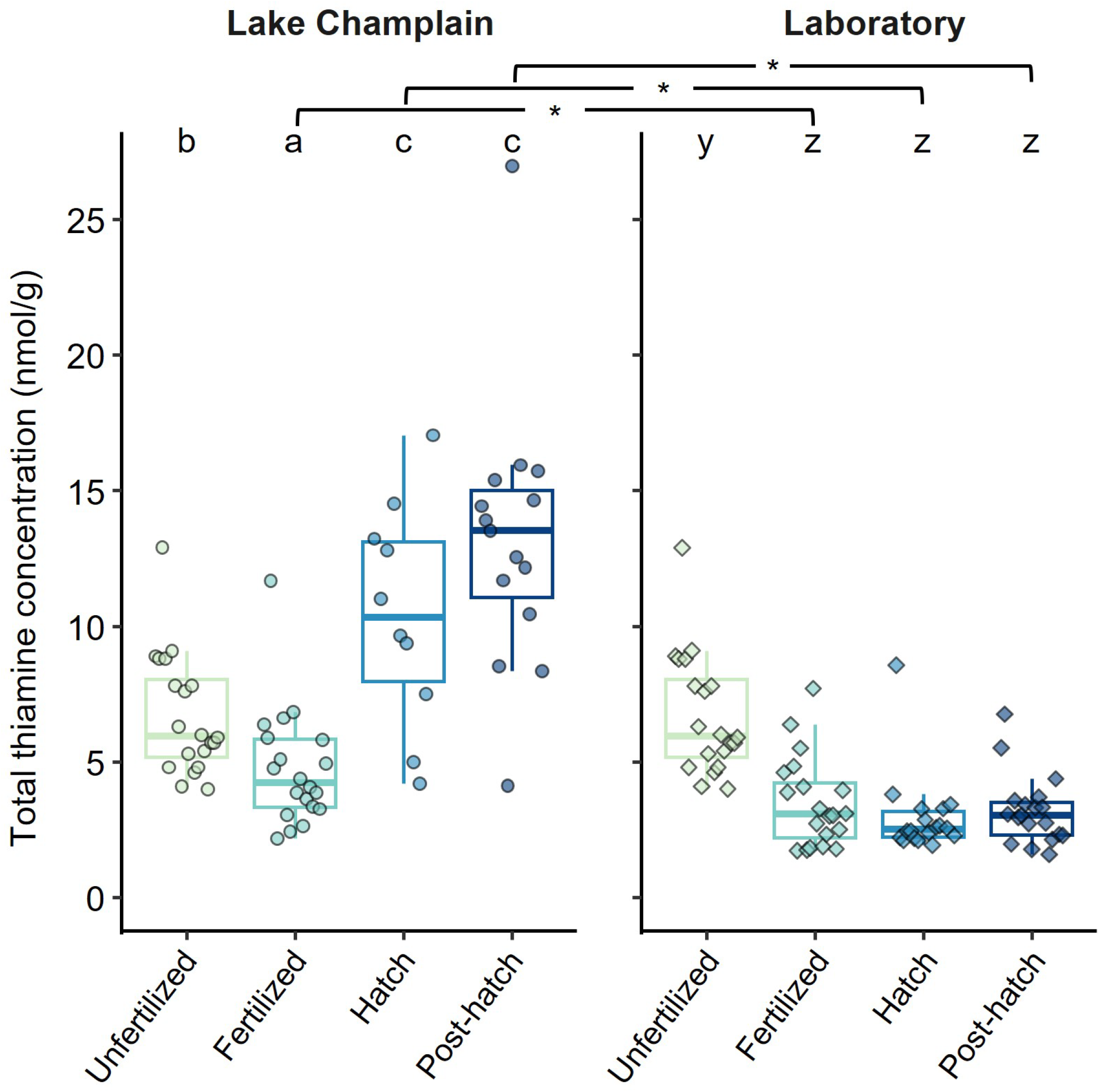
Variation in total thiamine concentrations of lake trout eggs or embryos sampled at four periods (unfertilized eggs, fertilized eggs, free embryos at hatch, and free embryos post-hatch) during development under lake (Lake Champlain) and laboratory conditions. Significant differences in concentrations across sampling periods within each site are indicated by differing letters and significant differences between sites during the same sampling period are indicated by an asterisk. Boxes represent the interquartile range with a median line and whiskers represent most extreme values within 1.5 times the interquartile range.

Temporal changes in tissue dry thiamine concentrations followed similar patterns as wet thiamine concentrations, including more pronounced increases throughout development for eggs and embryos reared in the lake compared to those reared in the lab (Figure 2). However, an initial decrease in thiamine concentrations following fertilization was not observed in the lake group (average decline = 2%; p = 0.999; estimated marginal means) and was less pronounced in the lab samples compared to the observed change in wet thiamine concentrations with an average decline of 37% following fertilization. Similar to wet thiamine concentrations, this decrease in the lab samples was statistically significant (p < 0.001; estimated marginal means). Following fertilization, lake-reared embryos had a significant increase in dry thiamine concentrations when sampled at hatch with an average increase of 136% compared to concentrations in fertilized eggs (p < 0.001; estimated marginal means). Average dry thiamine concentrations also increased significantly in the lake-reared embryos between hatch and the final sampling event (p = 0.001; estimated marginal means), increasing by 69%. In contrast, thiamine concentrations further declined by an additional 25% between fertilization and hatch in lab-reared embryos, although this change was not statistically significant (p = 0.172; estimated marginal means). Dry thiamine concentrations increased significantly, by 118%, in lab-reared embryos between hatch and the final sampling period (p < 0.001; estimated marginal means) and the final concentrations were statistically similar to initial concentrations (p = 0.999; estimated marginal means). Comparisons between lake- and lab-reared embryos at individual sampling periods showed similar thiamine concentrations following fertilization (average difference of 25%; p = 0.142; estimated marginal means), but significantly greater dry thiamine concentrations in lake-reared embryos at both the hatch (average difference of 121%; p < 0.001; estimated marginal means) and final (average difference of 104%; p < 0.001; estimated marginal means) sampling period (Figure 2).

**Figure 2.**
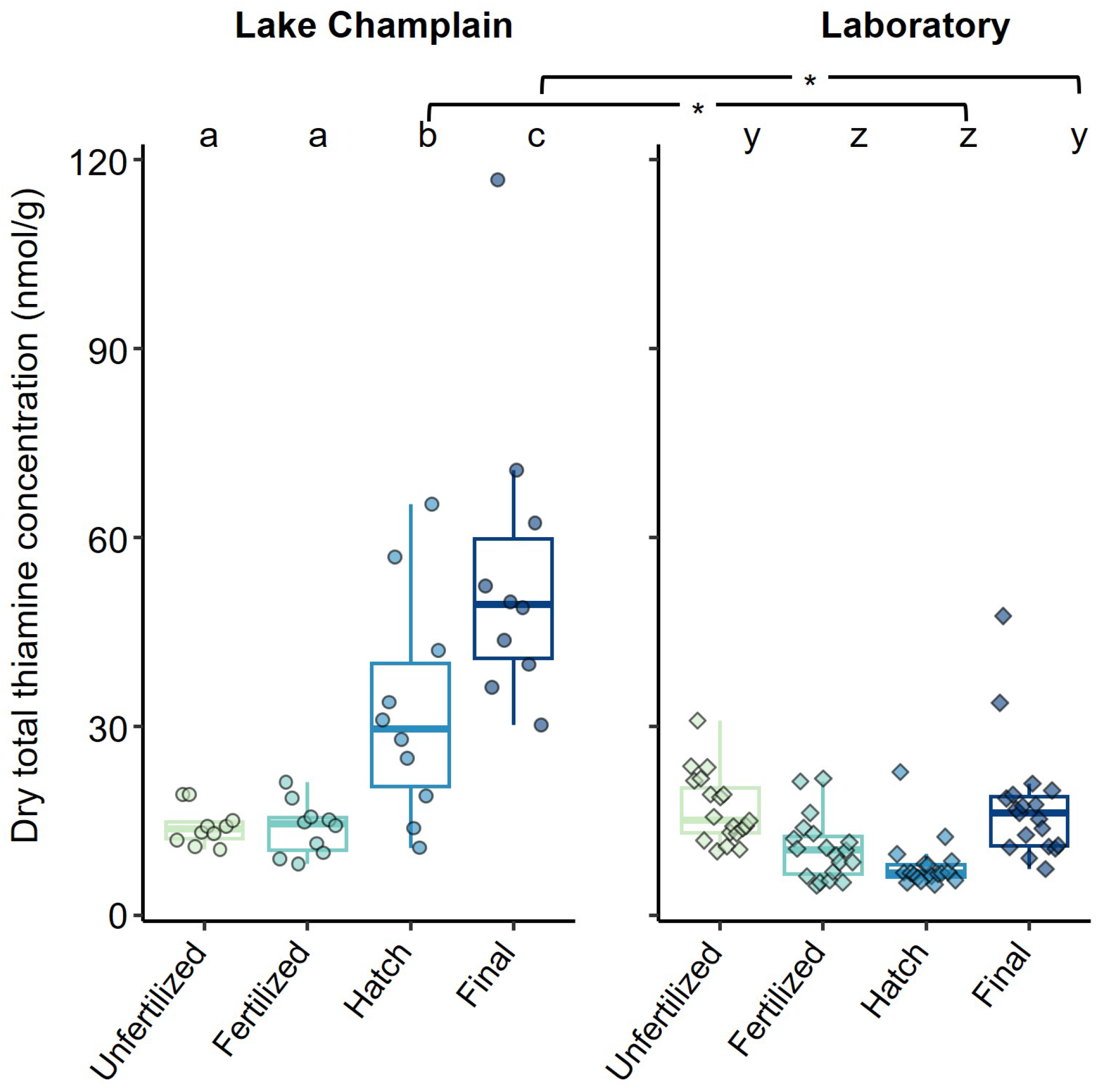
Variation in total thiamine concentrations from freeze dried lake trout eggs or embryos sampled at four periods unfertilized eggs, fertilized eggs, free embryos at hatch, and free embryos post-hatch) during development under lake (Lake Champlain) and laboratory conditions. Significant differences in concentrations across sampling periods within each site are indicated by differing letters and significant differences between sites during the same sampling period are indicated by an asterisk. Boxes represent the interquartile range with a median line and whiskers represent most extreme values within 1.5 times the interquartile range.

### Vitamer contributions

The percent contributions of thiamine vitamers in wet egg samples changed substantially among sampling periods for lake- and lab-reared eggs and embryos (Supplementary Figure 1). Thiamine pyrophosphate was the most abundant vitamer at all but one sampling period for both lake and lab groups. The one exception was fertilized eggs in the lab group, for which unphosphorylated thiamine was the most abundant vitamer. Trends in contributions of each thiamine vitamer throughout development were also largely similar between lake- and lab-reared groups. In both groups, percent contributions of thiamine pyrophosphate decreased between the unfertilized and fertilized egg sampling periods (61.1 ± 12.5% vs 48.1 ± 16.1% in lake samples and 35.4 ± 13.8% in lab samples) while unphosphorylated thiamine increased (27.2 ± 13.2% vs 41.3 ± 17.4 % in lake samples and 50.3 ± 17.1% in lab samples). Thiamine pyrophosphate then increased for free embryos sampled at hatch (89.1 ± 2.3% in lake samples and 79.0 ± 12.4% in lab samples), and further increased for free embryos at the final sampling period (95.8 ± 1.9% in lake samples and 92.4 ± 4.3% in lab samples). These large increases in thiamine pyrophosphate largely corresponded with decreases in thiamine monophosphate in individuals collected at hatch (3.4 ± 1.6% in lake samples and 11.4 ± 10.9% in lab samples) and the final sampling period (0.8 ± 0.6% in lake samples and 1.2 ± 1.4% in lab samples). Percent contributions of thiamine monophosphate were always below 25% among all samples and tended to decrease over time, except for a slight increase between the unfertilized and fertilized sampling periods for eggs in the lab group (Supplementary Figure 1).

The percent contribution of unphosphorylated thiamine in wet eggs and embryos differed between lake- and lab-reared groups, but the extent of the difference was dependent on sampling period (Figure 3). Fertilizing eggs with lab water led to significantly greater contributions of unphosphorylated thiamine in eggs compared to eggs fertilized with lake water (p < 0.001; GLMM). This difference was maintained when free embryos were sampled at hatch with significantly greater proportion of unphosphorylated thiamine in lab-reared free embryos compared to those reared in the lake (p = 0.024; GLMM). By the final sampling period post-hatch, contributions of unphosphorylated thiamine were similar between rearing groups (p = 0.116; GLMM) and were less than 5.5% among all samples. The same samples of unfertilized eggs were used for the initial sampling period for both lake and lab groups, and therefore no comparison was done for this stage.

**Figure 3.**
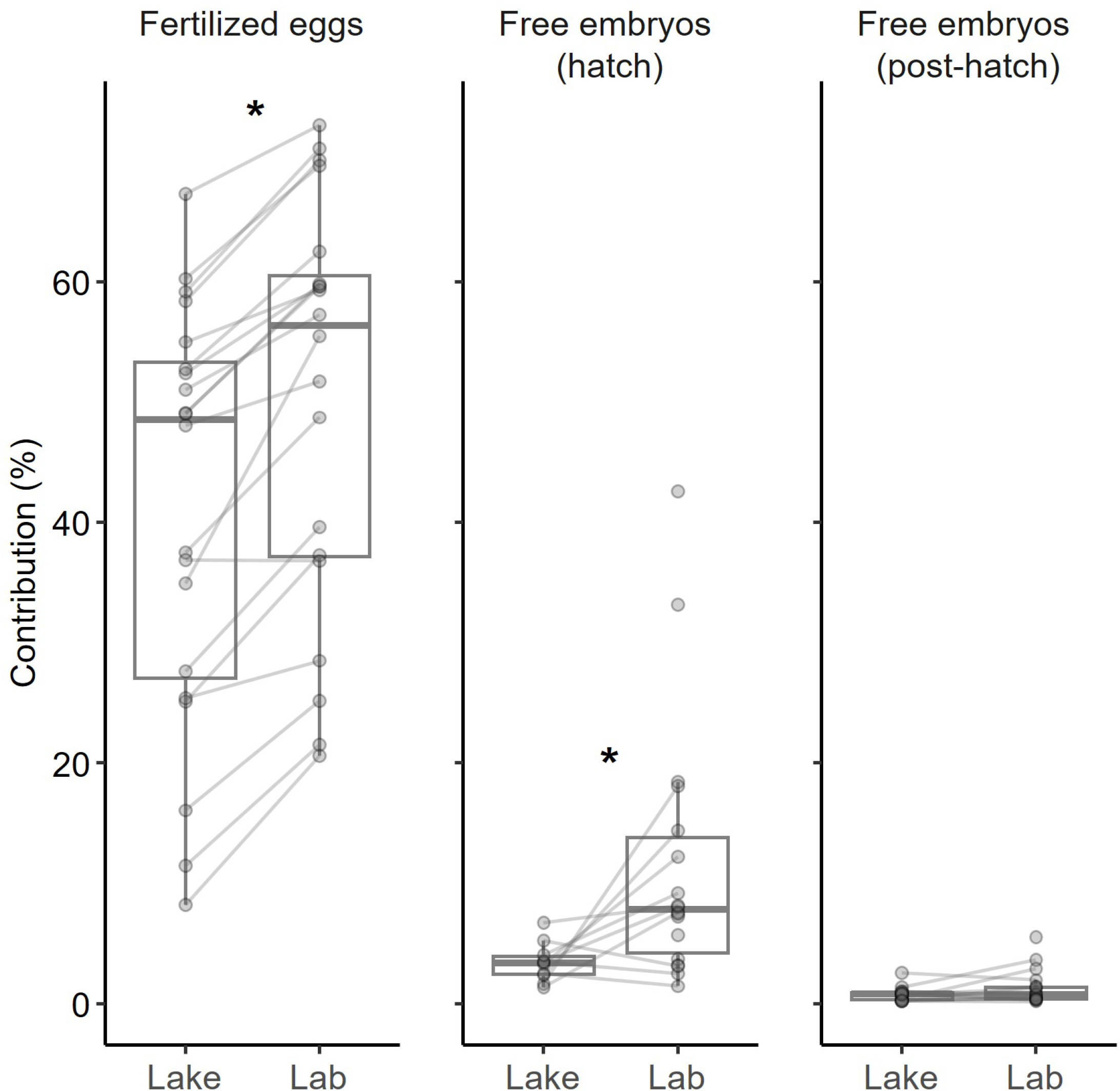
Percent of unphosphorylated thiamine in egg or embryo total thiamine concentrations compared between lake and laboratory rearing groups at three sampling periods (fertilized eggs, free embryos at hatch, and free embryos post-hatch). Datapoints within sampling periods are paired by family (i.e., same maternal parent), shown as connecting lines. Significant differences between groups by sampling period are indicated with an asterisk. Boxes represent the interquartile range with a median line and whiskers represent most extreme values within 1.5 times the interquartile range.

### Exogenous thiamine

We found measurable concentrations of dTRCs in water at each sampling period (fertilization, hatch, and post hatch), indicating active microbial thiamine biosynthesis and abiotic degradation was occurring in both the lake and lab environments during the approximately six months exposed to ambient water (Figure 4; Supplementary Table 2). However, there was substantial variability in compound concentrations among sampling periods and, in some cases, among rearing locations. The two thiamine-related precursors examined, the pyrimidine compound 4-amino-5-hydoxymethyl-2-methylpyrimidine (HMP) and the thiazole compound 5-(2-hydroxyethyl)-4-methyl-1,3-thiazole-2-carboxylic acid (cHET), showed different trends among sample locations and sampling periods. At fertilization, concentrations of HMP were high in water collected from both lake sites and were typically greater than the lab water samples. At the next two sampling periods, HMP concentrations were lower in water collected from both lake sites and were at the lower end of the range of concentrations observed for the corresponding lab samples. In contrast, cHET concentrations were relatively consistent across sampling periods with higher concentrations in water from both lake sites compared to the lab samples, except for one high concentration in water from the lab site observed during the initial sampling period. Thiamine degradation products including the thiazole compound 4-methyl-5-thiazoleethanol (HET) and the pyrimidine compound 4-amino-5-aminomethyl-2-methylpyrimidine (AmMP) had similar spatial and temporal trends. At fertilization, the highest concentrations for both compounds were in water samples from Blodgett Reef and lab samples were the lowest. HET concentrations in water for both lake sites dropped by the sampling at hatch and maintained similar concentrations until the final post-hatch sampling. AmMP concentrations dropped substantially in water from the Blodgett Reef site after fertilization then increased slightly by the final sampling, while concentrations in Gordon Landing samples remained similar across all three periods. HET and AmMP concentrations remained similar in lab samples among all three sampling intervals. HET concentrations were similar among water from all three sampling locations at both the hatch and final sampling intervals, but AmMP concentrations from lab samples were lower than in lake samples for all sampling periods. Lastly, unphosphorylated thiamine concentrations followed patterns similar to AmMP with the higher concentrations associated with water from both lake sites compared to the lab samples, which largely were consistent among all sampling periods. Relatively large variability was also observed for samples from both lake sites across and within sampling intervals, yet the range of concentrations for the two sites were typically similar to each other (Figure 4; Supplementary Table 3).

**Figure 4.**
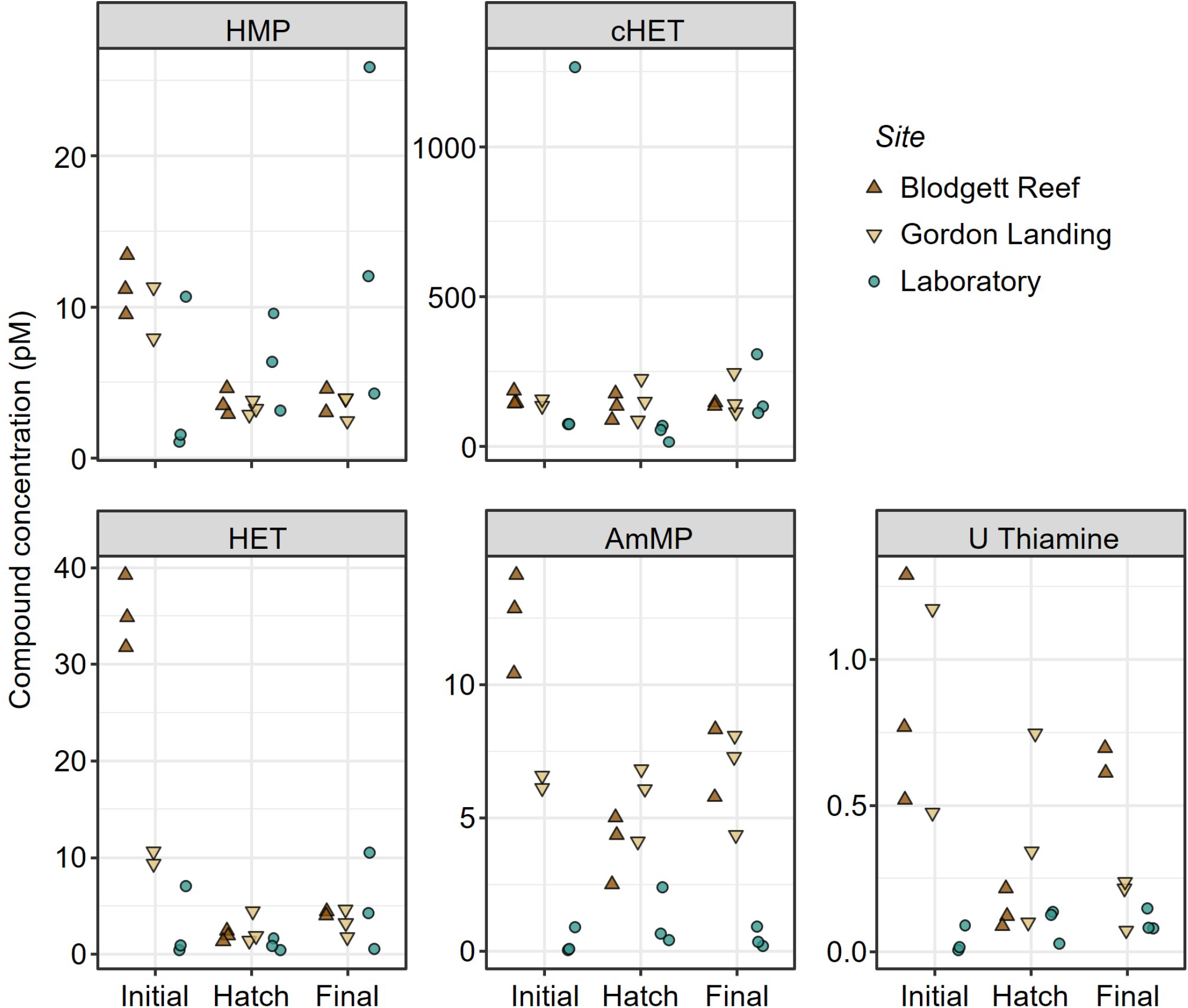
Concentrations (pM) of dissolved thiamine related compounds (dTRC) including the thiamine precursors 4-amino-5-hydoxymethyl-2-methylpyrimidine (HMP) and 5-(2-hydroxyethyl)-4-methyl-1,3-thiazole-2-carboxylic acid (cHET); the thiamine degradation products 4-methyl-5-thiazoleethanol (HET) and 4-amino-5-aminomethyl-2-methylpyrimidine (AmMP); and unphosphorylated thiamine (U Thiamine). Compounds are grouped by sampling location with the two lake sites indicated by triangles and the laboratory site is represented by circles. Compounds are further separated by sampling period including the initial sampling at fertilization (Initial), sampling when eggs hatched (Hatch), and the final sampling post-hatch (Final). Note the change in y-axis scale among compounds.

Of the 296 free-embryo digestive tracts examined, only three contained any exogenous content, and those embryos were from three separate families. Thus, overall foraging success was only 1%. Based on the proportion of fish examined in these families, foraging success ranged from 4 to 11% (Table 1). Initial unfertilized egg thiamine concentrations for those three families were comparable to concentrations from other families (Table 1).

**Table 1.** Free embryo foraging success (%) for individual lake trout families based on presence of exogenous contents in the digestive tract when sampled from Lake Champlain (Gordon Landing) approximately five weeks post-hatch. The number of free embryos analyzed for each family (embryo count) and the corresponding total thiamine concentrations (nmol/g) from unfertilized eggs are also included. One family was missing an identification tag when sampled post-hatch and thus could not be associated with an initial thiamine concentration.

| Initial thiamine | Embryo count | Foraging embryos | Foraging success (%) |
| --- | --- | --- | --- |
| 9.07 | 20 | 1 | 5 |
| 7.28 | 10 | 1 | 10 |
| 4.13 | 25 | 1 | 4 |
| 12.90 | 15 | 0 | 0 |
| 8.88 | 15 | 0 | 0 |
| 7.81 | 18 | 0 | 0 |
| 7.75 | 18 | 0 | 0 |
| 7.65 | 24 | 0 | 0 |
| 6.26 | 22 | 0 | 0 |
| 5.99 | 12 | 0 | 0 |
| 5.88 | 12 | 0 | 0 |
| 5.75 | 15 | 0 | 0 |
| 5.42 | 15 | 0 | 0 |
| 4.84 | 14 | 0 | 0 |
| 4.80 | 15 | 0 | 0 |
| 4.60 | 20 | 0 | 0 |
| 4.05 | 17 | 0 | 0 |
| - | 9 | 0 | 0 |

## Discussion

Our results clearly demonstrate that developing lake trout embryos can acquire thiamine and increase reserves of this vitamin during development. While we were unable to determine whether these increases were sufficient to alleviate TDC and increase offspring survival, these results raise the question of whether previous studies that evaluate offspring mortality associated with TDC in hatchery settings represent offspring mortality rates in wild populations. The observed increases in thiamine concentrations during development are most likely associated with uptake of dTRCs from the surrounding environment, as most digestive tracts that were examined contained no food.

The difference in thiamine concentrations between the lab and lake environments indicates that rates of thiamine acquisition can vary depending on rearing environment and developmental period. The developmental stage associated with the greatest thiamine acquisition also differed between rearing environments. Most thiamine acquisition for lake-reared embryos occurred between fertilization and hatching whereas thiamine concentrations only increased post-hatch in the lab-reared group. However, the fastest rate of thiamine acquisition for the lake-reared embryos also occurred post-hatch considering the increase occurred in a matter of weeks compared to months between fertilization and hatching. The higher concentrations of various dTRCs at fertilization in the lake compared to the lab may be responsible for this difference in timing of thiamine acquisition. However, this hypothesis is confounded by the fact that the timing of sampling at hatch differed between the two rearing environments, causing the sampling periods to represent slightly different development periods. In the lab, newly-hatched embryos were sampled daily, so all embryos were sampled within 24 hrs of hatching thereby limiting post-hatch exposure to dTRCs. In contrast, embryos reared in the lake likely hatched multiple days prior to sampling considering nearly all live embryos had hatched by the time sampling occurred and the hatching period can last over ten days among eggs fertilized on the same date^[29]^. Thus, exposure to dTRCs between hatching and sampling for individuals in the lake-reared group may have influenced the significant increase in thiamine concentrations observed between the fertilization and hatch periods. Variability in exposure time to ambient dissolved thiamine between hatching and sampling among lake-reared embryos may also have contributed to the relatively high amount of variation in thiamine concentrations among embryos at that sampling period compared to the amount of variation among either other lake samples (i.e., sampled at fertilization or final sampling post-hatch) or any lab-reared samples. These differences in thiamine acquisition between lake- and lab-reared embryos are also similar to differences observed previously in TDC-related offspring mortality rates between siblings reared in two separate facilities^[30]^, further indicating that thiamine acquisition is possible and site-dependent.

Although wet thiamine concentrations declined between initial unfertilized egg collection and sampling after fertilization, this effect was likely due to dilution; ambient water is absorbed by eggs during water hardening and can increase egg volume by approximately 20%^[31]^. Taking that dilution factor into consideration, the significantly greater wet thiamine concentrations in lake-reared embryos compared to lab-reared embryos following fertilization suggests lake-reared embryos obtained more ambient thiamine during the fertilization and water hardening processes. This conclusion is also supported by the smaller decrease in wet and dry thiamine concentrations following fertilization for lake-reared embryos compared to lab-reared embryos. Differences in thiamine acquisition during fertilization also changed thiamine composition between the sampling groups. Interestingly, while no thiamine appeared to be acquired by the eggs fertilized with lab water, the percentage of unphosphorylated thiamine in those eggs increased substantially. Eggs fertilized with lake water also experienced an increase in the percent contribution of unphosphorylated thiamine, but to a significantly lower extent. Unphosphorylated thiamine is the mobile form of thiamine and is typically the most abundant vitamer in healthy salmonine eggs serving as a reserve of the vitamin^[21]^. As the mobile form of thiamine, unphosphorylated thiamine may be more readily available dissolved in ambient water than other vitamers, potentially explaining the greater contributions of unphosphorylated thiamine observed in eggs following fertilization. It is not clear if or how these changes in vitamer contributions and thiamine concentrations following fertilization may influence TDC, but considering increases in total thiamine concentrations during this period were minimal and relative abundance of vitamers were similar between rearing groups by the final sampling period, we expect any health effects would be minimal.

Thiamine acquisition in natural settings differs from artificial treatments both in the timing and intensity (i.e., concentration) of exposure to thiamine. Thiamine concentrations in natural waters are substantially lower than in water used to treat embryos in culture, but uptake could occur over a much longer period of exposure, five to six months during embryo development within the egg. The comparison, therefore, is between uptake during a short period of high thiamine availability, which is an acute exposure, and during a long period of low thiamine availability, or chronic exposure. Acute exposure (i.e., 1 hr) during water hardening using egg baths have been shown to be ineffective when supplementation of thiamine hydrochloride is below 500 mg/L (equivalent to 1.5*10^9^ pM/L)^[32]^, which is much greater than any natural dTRC concentrations we observed (≤ 1300 pM/L) or reported in other systems^[25,33,34]^. Our results, however, indicate that chronic exposure (i.e., months) to low concentrations of dTRCs in natural environments allows thiamine concentrations to increase gradually over time. The exact timing and rate of thiamine acquisition during chronic exposure is unknown, but it is unlikely to be consistent throughout development. Concentrations of dTRCs can be highly variable over time and the ability of eggs or developing embryos to take in ambient thiamine may also change across development stages. Most thiamine treatments occur shortly after fertilization during water hardening^[22,23,32]^ or after hatching^[23,24]^, presumably when acquisition is most efficient. Thiamine acquisition may be reduced during the incubation period following water hardening due to the limited rate of small-molecule diffusion across the chorion and underlying membranes^[35]^.

Natural thiamine sources include dTRCs and prey (i.e., zooplankton), and studies have suggested that both of these sources may mitigate TDC if available^[25,26]^. We observed comparable concentrations of most dTRCs between the lab and both lake sites, although some unphosphorylated thiamine and AmMP concentrations were higher at the Gordon Landing site for the hatch and post-hatch sampling periods when thiamine acquisition seemed greatest. In contrast, HMP concentrations tended to be higher in the lab samples compared to lake samples at hatch and the final sampling period post-hatch. However, low dissolved concentrations of biologically important compounds can indicate high rates of biological uptake rather than low availability^[33]^. It remains unclear whether these differences between lab and lake dTRC concentrations would be sufficient to cause the observed differences in thiamine acquisition between the lake- and lab-reared embryos. As an alternative hypothesis, altered embryo gut microbiome associated with the different rearing environments^[36]^ may cause lower thiamine assimilation efficiency for fish reared in lab. Additionally, bacteria in the gastrointestinal tract can produce thiamine, which can then be absorbed by the host^[37]^, so differences in the bacterial community may influence thiamine obtained by the host. No research to date has explored either of these microbial effects in fishes and the potential effects on the extent of TDC. Interestingly, we also noticed substantial temporal and spatial variability in dTRC concentrations between the two lake sites, indicating thiamine availability within Lake Champlain were site-dependent similar to other systems^[25,34]^. However, all compounds from both lake sites had concentrations that fell within the range of concentrations reported from other freshwater systems (Sacramento River watershed, California and Klamath Lake), though maximum values reported in the Sacramento River watershed were substantially greater than maximum values observed in Lake Champlain^[25]^. The concentrations for dTRCs from the lab samples were also within the range of concentrations observed in other natural systems, but many were towards the lower end of previously reported concentrations compared to Lake Champlain samples^[25,34]^.

Variability in prey availability and foraging rates may also lead to different thiamine acquisition spatially and temporally. Zooplankton likely have sufficient thiamine concentrations to increase embryo thiamine concentrations^[27]^, and lake trout in Lake Champlain have been observed feeding as soon as two weeks after hatching^[26]^. Interestingly, stomachs in our study were largely empty which may result from temporal variation in prey availability or differences in how embryos were captured. Zooplankton availability can be variable across years depending on lake conditions^[38]^, so it is possible that zooplankton abundance was low during spring of our study. Zooplankton abundance can also vary spatially throughout the substrate and water column. Previous assessments of free embryo lake trout foraging used fry emergence traps that collect embryos during diurnal vertical migrations from the substrate and thus are more likely to interact with zooplankton. Fry trap capture bottles also allow free embryos to swim freely in ambient water that contains zooplankton. In contrast, the egg bags used in this study trapped embryos in the reef substrate where zooplankton abundance and freedom of movement by embryos may have been lower. Thus, unconstrained embryos are likely to have higher feeding rates, as observed previously^[22]^, and may have larger increases in thiamine than we observed.

The comparisons between lab and lake reared embryos in this study were imperfect due to differences in water temperatures between the lab and lake sites, thus leading to different development rates and exposure times to any ambient nutrients for a given development stage. As a result, the period between fertilization and hatch was much shorter for lab-reared embryos compared to those reared in the lake, while the period between hatch and the final sampling was much longer for lab-reared embryos compared to lake-reared embryos. Given the rate of thiamine acquisition was greatest after hatching, this difference in development period durations would likely provide lab-reared individuals more time exposed to ambient thiamine during a period when acquisition is most efficient compared to lake-reared individuals. That longer exposure may have contributed to the increased dry thiamine concentrations observed for lab-reared embryos between hatch and the final sampling period, yet this increase was still lower than the increase observed for lake-reared embryos during that same development stage despite the longer exposure time. Thus, if development rates were comparable across lake- and lab-reared fish with consistent exposure times to thiamine, particularly after hatch, differences in thiamine acquisition post-hatch could have been even greater than what was observed in this study.

A major limitation of this study was the inability to assess free-embryo behaviors and survival relative to their rearing environment and thiamine acquired. Previous research has confirmed that thiamine acquisition during development can reduce mortality associated with TDC^[22–24]^, but it is not clear whether the quantity of thiamine acquired by embryos in this study would have been sufficient to offset health issues associated with TDC. Initial thiamine concentrations in unfertilized eggs were likely sufficient for most embryos to avoid direct mortality caused by TDC^[21]^. Secondary effects associated with higher thiamine concentrations^[5]^ would likely still occur, especially for lab-reared embryos that experienced little to no increase in thiamine concentrations. These secondary effects can also restrict free embryo activity and foraging success^[5]^, which may have contributed to the low incidence of foraging we observed. Additionally, if most thiamine acquisition occurs after hatching, long-term consequences of TDC may still occur, including reduced growth and impaired immune response^[39,40]^. However, it is important to note that offspring survival has been sufficient to produce the first year classes of wild recruits since stocking began in 1972, and ultimately restore a wild population of lake trout to Lake Champlain^[19]^ (M. Murphy, VTFWD, pers. comm.) despite low thiamine concentrations in adults^[26]^. Thus, it is possible that ambient thiamine in Lake Champlain may be reducing TDC and contributing to the observed recruitment. However, other factors such as lower thiamine requirements for the strain of lake trout in Lake Champlain (i.e., Seneca strain) may also contribute to increased recruitment^[41]^.

This study is the first to document thiamine acquisition by developing salmonine embryos in a natural environment, including substantial increases in embryo thiamine concentrations. The observed increases in thiamine concentrations occurred without any thiamine supplementation, indicating thiamine was obtained from natural sources. While it remains to be determined if these increases in thiamine concentrations can effectively offset effects of TDC, spatial and temporal variability in ambient thiamine may contribute to differences in the extent of TDC and therefore natural recruitment observed among systems or over time. Future research looking into thiamine concentrations in embryos from healthy populations would help to evaluate the implications of increased embryo thiamine concentrations observed in this study and the potential for reduced TDC in natural populations. Increased thiamine in embryos developing in natural environments relative to siblings reared in artificial (i.e., hatchery) environments suggests that the limitations on recruitment resulting from TDC may be lower than previously expected based on mortality estimates developed from lab-based studies.

## Methods

### Fish collection and embryo fertilization

Lake trout gametes were collected from mature adults captured during the spawning season in Lake Champlain (44.6881, -73.3487) using trap nets deployed by the Vermont Fish and Wildlife Department during fall 2021. Eggs were stripped from 20 females, and eggs from each female were fertilized using milt pooled from three males. Each family was split into two groups for lake rearing and lab rearing. Water used to mix gametes for fertilization was obtained from the corresponding environment where the eggs would be reared (i.e., lab or lake). Milt was introduced to batches of approximately 500 eggs, then water was added to cover the mixture for one min to allow fertilization. Eggs were then rinsed with their respective water to remove excess milt and resubmerged in 1 L of fresh water for water hardening. After water hardening for two hours, a sample of approximately 100 eggs was collected from each family for thiamine analysis and remaining eggs were again subdivided into an additional two groups (approximately 200 eggs each) corresponding to the two later sampling periods at hatch and post-hatch.

### Embryo rearing and collection

Eggs in the lake-reared group were reared in egg bags^[42]^ deployed in the rocky substrate at the base of a breakwall at Gordon Landing in Lake Champlain, a site that has produced large numbers of free embryos since studies began there in 2000^[43]^. Egg bags were buried in cavities created by scuba divers in the breakwall substrate and backfilled with rocks to provide substrate for embryos and to help separate eggs to minimize fungal growth. Egg bags were arranged in a predetermined and random order along the reef. On November 10^th^, 2021, approximately 200 fertilized eggs were added to each egg bag by scuba divers and bags were then covered to prevent escapement after hatch and limit predation on embryos; bags were numbered corresponding to family and sampling period. Approximate hatch date was estimated based on daily temperature recordings provided by a US Geological Survey monitoring station located on Lake Champlain. Temperatures were used to calculate cumulative degree days and predict approximate hatch dates (approximately 400-450 cumulative degree days when reared at about 4.4 °C^[44]^) for the first sampling of reared embryos. Embryos were sampled on April 5^th^, 2022 at approximately 480 cumulative degree days. The second sampling of reared embryos occurred approximately five weeks later on May 13^th^, 2022 at approximately 630 cumulative degree days. All embryos sampled were frozen immediately on dry ice and stored at -80 °C until processing.

Embryos in the lab group were reared at the Rubenstein Ecosystem Science Laboratory in Burlington, Vermont (United States). Eggs were reared in vertical incubators (i.e., Heath trays) with families separated in individual polyvinyl chloride containers with a mesh bottom and lid to allow water flow. The hatchery system used recirculating dechlorinated municipal water that was chilled to 4.8 °C (daily average and one standard deviation = 4.8 ± 0.3 °C). Each hatchery system included two 38 L aquarium filters with activated carbon mesh filter cartridges. Water temperatures were recorded daily for each tray and water quality (ammonia, nitrate, and nitrate concentrations) was assessed biweekly for each system. Embryos started hatching January 24^th^, 2022, with peak hatch approximately February 1^st^, 2022 (approximately 400 cumulative degree days). Embryos were observed daily throughout the rearing period to remove dead individuals and identify hatched individuals. Once hatched, free embryos were either frozen on dry ice for thiamine analyses or transferred to 6.7 L aquaria with one family per aquarium. The aquaria were maintained on the same water system as incubating eggs until the final collection, which occurred on April 21^st^, 2022 at approximately 785 cumulative degree days.

A subsample of lake-reared free embryos from the final sampling post-hatch was held separately to count the number of feeding individuals, identified by presence of zooplankton in stomachs. These embryos were preserved in 70% ethanol immediately after capture and stored in that solution until processing. Stomach contents were analyzed for 17 of the 20 lake-reared families; insufficient live individuals were available from the other three families. Between nine and 25 individuals were sampled per family, with a total of 296 free-embryo stomachs dissected (Table 1). Presence of exogenous materials in the digestive tract was assessed visually after removing the entire tract from individual embryos.

Lake trout collections from Lake Champlain were conducted under a Vermont State Scientific Collection permit (authorization # S-2020 EM). The protocol for this experiment and rearing of embryos was approved by the University of Vermont Institutional Animal Care and Use Committee (IACUC; protocol # PROTO202100057) in accordance with the requirements of the Office of Laboratory Animal Welfare, National Institutes of Health (Animal Welfare Assurance #A3301-01), and following ARRIVE guidelines^[45]^.

### Water sampling

Water samples were collected from two lake reefs and the hatchery system at three periods during the study period: fertilization, at hatch, and post hatch. The two lake reefs were the Gordon Landing breakwall and a submerged breakwall ruin, Blodgett Reef, located approximately 25 km south of the Gordon Landing breakwall where lake trout free embryos have been sampled in the past (unpublished data). Two 2-L samples of reef and dechlorinated city water were collected at each sampling period for thiamine analysis. Samples of reef water were collected from interstices by scuba divers or using a hand-operated bilge pump with a weighted hose dropped to the substrate surface. All water samples were acidified using 2 mL of 1 M HCl and filtered with acid-washed polycarbonate membrane filters (0.2 μm), then frozen at -20 ºC until further processing.

### Thiamine quantification

Thiamine vitamers were extracted from egg and embryo tissue in duplicate^[28]^. Briefly, approximately 1 g of tissue was homogenized for one minute in a chilled 1.5 mL solution of 2% trichloroacetic acid. Samples were then boiled at 100 °C for 15 min followed by 15 min of centrifugation at 14,000 g. The supernatant was removed and washed four times with an ethyl acetate:hexane (3:2 ratio) solution. A combination of 1.2N NaOH and 0.1% potassium ferricyanide were added to each sample and samples were then passed through a polytetrafluoroethylene-membrane filter with 0.2 µm pore size. Concentrations of unphosphorylated thiamine, thiamine mononitrate, and thiamine pyrophosphate were each quantified from filtered samples using high performance liquid chromatography (HPLC)^[22]^. Concentrations of each vitamer were determined by comparing peak area to a six-point standard curve with known concentrations of thiamine that was generated at the start of each HPLC run. Vitamer concentrations for each sample were summed to determine the total thiamine concentration. An average thiamine concentration was then calculated from the two replicates for each sample. Thiamine was extracted and quantified at State University of New York Brockport.

Thiamine concentrations in dehydrated samples (i.e., dry thiamine concentration) were estimated by adjusting tissue weight from thiamine extraction to account for water content. Water content was determined by freeze drying approximately 1 g of remaining tissue for each sample and calculating the change in mass following dehydration. Thiamine concentrations from hydrated samples were then converted to dry thiamine based on the estimated dry weight of the initial (hydrated) sample. Samples were dried for approximately 20 hr at -20 °C.

Dissolved thiamine-related compounds were extracted from preserved lakewater using solid phase extraction followed by liquid chromatography mass spectrometry (LCMS) to measure dTRC concentrations as described previously^25^. The dTRCs included unphosphorylated thiamine, the thiamine precursors pyrimidine compound 4-amino-5-hydoxymethyl-2-methylpyrimidine (HMP) and thiazole compound 5-(2-hydroxyethyl)-4-methyl-1,3-thiazole-2-carboxylic acid (cHET), and the thiamine degradation products thiazole compound 4-methyl-5-thiazoleethanol (HET) and pyrimidine compound 4-amino-5-aminomethyl-2-methylpyrimidine (AmMP). Compound concentrations for each extracted sample were analyzed in triplicate and were processed in a randomized order. An internal standard (^13^C-labeled thiamine) was used for quantification and to compensate for matrix effects. The LCMS analysis was conducted at the Oregon State University Mass Spectrometry Core Facility.

### Data analyses

The extent of thiamine acquisition was assessed by comparing thiamine within families across sampling periods. Variation in thiamine concentrations among sampling periods and between sampling sites was assessed for wet and dry embryo samples using generalized linear mixed effects model (GLMM). A separate model was run for each sample type (wet or dry) using the `glmmTMB` function from the *glmmTMB* R package^[46]^. Both GLMMs included site (lake or lab) and stage (unfertilized, fertilized, hatch, and post-hatch) as fixed effects and a family identifier as a random effect. The GLMMs were fit to a gamma distribution and included a log link function. Post-hoc analyses within sample types were conducted for pairwise comparisons between stages and sites based on estimated marginal means with a Bonferroni correction for multiple comparisons using the `emmeans` function from the *emmeans* R package^[47]^. Contributions of each thiamine vitamer towards total thiamine concentrations were assessed as percentage of total thiamine at each sampling period. Differences in the percent contribution of unphosphorylated thiamine between lake and lab reared groups were assessed using GLMMs with the `glmmTMB` function for each sampling period with site as the only fixed effect and family as a random effect. Percent contributions of unphosphorylated thiamine were normally distributed at the fertilized sampling period and therefore the GLMM used for those data was fit to a gaussian distribution. However, data were non-normal for sampling at hatch and final sampling post-hatch, so the GLMMs used for those sampling periods used the beta family distributions with a logit link function. Concentrations for each dTRC were averaged among sample triplicates for each 1 L sample, resulting in up to three values for each compound by sampling period (fertilization, hatch, and post-hatch); only two samples were successfully processed for the initial sampling period at Gordon Landing and the final sampling period at Blodgett Reef. Due to limited sample size, comparisons of dTRC concentrations among sample sites (two lake and one lab site) and periods (fertilization, at hatch, and post-hatch) were not evaluated for statistical significance. All analyses were conducted using R^[48]^ with figures generated using the `ggplot` function in the *ggplot2* R package^[49]^.

## Acknowledgments

This project could not be completed without the assistance of numerous collaborators and assistants in the field and lab. We thank the Vermont Fish and Wildlife Department for providing lake trout, crew of the R/V Melosira at the University of Vermont and divers from Vermont Diving for assistance deploying and recovering equipment and sample, members of the Rubenstein Ecosystem Science Laboratory for and manuscript review, and students from the Rinchard Lab for assistance with biochemical analyses in egg and embryos.

## Author contributions

MHF contributed to conceptualization, data curation, formal analysis, investigation, methodology, project administration, resources, supervision, visualization, writing – original draft preparation, and writing – review and editing. CC contributed to data curation, formal analysis, investigation, and project administration. CPS contributed to data curation, methodology, project administration, and writing – review and editing. GSJ contributed to data curation, project administration, and writing – review and editing. JEM contributed to conceptualization, data curation, investigation, methodology, project administration, resources, supervision, and writing – review and editing. JR contributed to conceptualization, data curation, investigation, methodology, project administration, resources, supervision, and writing – review and editing.

## Data availability statement

The datasets generated and analysed during the current study are available from the corresponding author on reasonable request.

