## Supplementary Materials for "Thiamine availability and acquisition differ between natural and controlled environments"

**Supplementary Information**

Supplementary Table 1. Thiamine vitamer and total thiamine (TTH) concentrations for unfertilized eggs collected from 20 females sampled during fall spawning at Gordon Landing in Lake Champlain. Vitamers include unphosphorylated thiamine (TH), thiamine monophosphate (TMP), and thiamine pyrophosphate (TPP).

| Family | TH | TMP | TPP | TTH |
| --- | --- | --- | --- | --- |
| 1 | 3.68 | 0.85 | 4.35 | 8.88 |
| 2 | 3.10 | 0.84 | 4.84 | 8.77 |
| 3 | 6.58 | 1.18 | 5.14 | 12.90 |
| 4 | 2.39 | 0.78 | 4.11 | 7.28 |
| 5 | 0.46 | 0.71 | 3.67 | 4.84 |
| 6 | 3.15 | 0.93 | 4.99 | 9.07 |
| 7 | 3.15 | 0.96 | 3.70 | 7.81 |
| 8 | 0.49 | 0.69 | 5.08 | 6.26 |
| 9 | 0.34 | 0.56 | 3.23 | 4.13 |
| 10 | 3.33 | 0.89 | 3.42 | 7.65 |
| 11 | 1.66 | 0.78 | 2.89 | 5.33 |
| 12 | 2.78 | 0.83 | 4.14 | 7.75 |
| 13 | 0.73 | 0.51 | 3.37 | 4.60 |
| 14 | 0.79 | 0.66 | 3.35 | 4.80 |
| 15 | 1.58 | 0.79 | 3.62 | 5.99 |
| 16 | 1.30 | 0.68 | 3.44 | 5.42 |
| 17 | 0.50 | 0.48 | 3.07 | 4.05 |
| 18 | 0.97 | 0.73 | 3.97 | 5.67 |
| 19 | 2.37 | 0.70 | 2.68 | 5.75 |
| 20 | 1.18 | 0.73 | 3.96 | 5.88 |

Supplementary Table 2. Summary concentrations (pm) for dissolved thiamine related compounds (dTRCs) including the thiamine precursors pyrimidine compound 4-amino-5-hydoxymethyl-2-methylpyrimidine (HMP) and thiazole compound 5-(2-hydroxyethyl)-4-methyl-1,3-thiazole-2-carboxylic acid (cHET), the thiamine degradation products thiazole compound 4-methyl-5-thiazoleethanol (HET) and pyrimidine compound 4-amino-5-aminomethyl-2-methylpyrimidine (AmMP), and unphosphorylated thiamine (U Thiamine). Concentrations are reported as averages among triplicates with minimum and maximum values included in parentheses. Samples are divided by sampling locations including two sites in Lake Champlain, Blodgett Reef and Gordon Landing, and one laboratory (lab) site. Samples are further divided into three sampling periods including the initial sampling which occurred at fertilization, sampling at hatch, and the final sampling post-hatch.

|  |  | HMP | cHET | HET | AmMP | U Thiamine |
| --- | --- | --- | --- | --- | --- | --- |
| **Blodgett**  **Reef** | Initial | 11.4 (9.5, 13.5) | 157.8 (142.7, 186.0) | 35.3 (31.8, 39.3) | 12.5 (10.4, 14.1) | 0.9 (0.5, 1.3) |
|  | Hatch | 3.7 (2.9, 4.6) | 133.6 (89.0, 177.0) | 1.9 (1.3, 2.4) | 4.0 (2.5, 5.0) | 0.1 (0.1, 0.2) |
|  | Final | 3.8 (3.0, 4.6) | 140.5 (134.7, 146.3) | 4.2 (4.0, 4.4) | 7.1 (5.8, 8.3) | 0.7 (0.6, 0.7) |
| **Gordon Landing** | Initial | 9.6 (7.9, 11.3) | 146.7 (135.4, 158.0) | 10.0 (9.4, 10.6) | 6.4 (6.1, 6.6) | 0.8 (0.5, 1.2) |
|  | Hatch | 3.3 (2.9, 3.8) | 154.1 (87.0, 225.3) | 2.6 (1.4, 4.4) | 5.7 (4.1, 6.8) | 0.4 (0.1, 0.7) |
|  | Final | 3.5 (2.5, 4.0) | 166.7 (113.3, 245.7) | 3.2 (1.8, 4.6) | 6.6 (4.4, 8.1) | 0.2 (0.1, 0.2) |
| **Lab** | Initial | 4.4 (1.1, 10.7) | 472.5 (75.3, 1,266.7) | 2.8 (0.4, 7.1) | 0.3 (0.0, 0.9) | 0.0 (0.0, 0.1) |
|  | Hatch | 6.4 (3.2, 9.6) | 46.3 (15.0, 68.4) | 1.0 (0.4, 1.6) | 1.2 (0.4, 2.4) | 0.1 (0.0, 0.1) |
|  | Final | 14.1 (4.3, 25.9) | 184.4 (111.3, 308.0) | 5.1 (0.6, 10.5) | 0.5 (0.2, 0.9) | 0.1 (0.1, 0.1) |

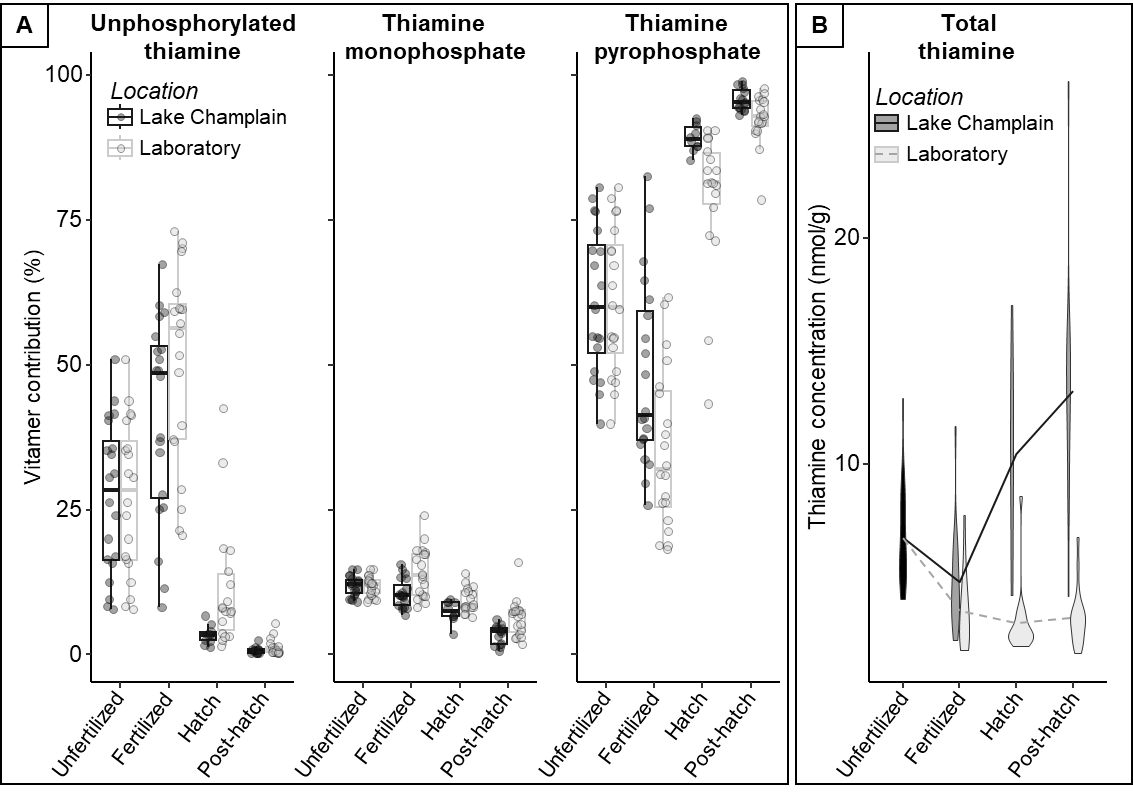
Supplementary Figure 1. (A) Percent contributions of thiamine vitamers (unphosphorylated thiamine, thiamine monophosphate, and thiamine pyrophosphate) towards total egg thiamine concentrations at four sampling periods (unfertilized, fertilized, at hatch, and final sampling post-hatch) separated by collection location (Lake Champlain or laboratory). (B) Variability in total thiamine concentrations across sampling periods for the lake and laboratory groups are also shown for reference. Boxes represent the interquartile range with a median line and whiskers represent most extreme values within 1.5 times the interquartile range.
